# A cell-intrinsic steroidogenic checkpoint constrains female Th2 effector output independently of gonadal hormones

**DOI:** 10.64898/2026.05.13.724806

**Authors:** Jhuma Pramanik, Qiuchen Zhao, Soura Chakraborty, Chengxuan Xie, Bidesh Mahata

## Abstract

T helper 2 (Th2) lymphocytes drive type-2 immunity and allergic diseases that disproportionately affect females, yet the mechanisms underlying this sexual dimorphism remain poorly understood. Current models attribute sex-biased immune differences to circulating gonadal hormones and to sex chromosome composition, leaving unresolved whether immune cells possess cell-autonomous mechanisms acting independently of both. Here we show that female and male Th2 cells differentiated under identical, hormone-free conditions are transcriptionally distinct but functionally equivalent: female cells carry a heightened inflammatory and interferon-response programme, yet produce IL-13 and IL-4 at rates indistinguishable from male cells, with only a modest baseline difference in proliferative capacity. Using T cell-specific deletion of *Cyp11a1* that encodes the first and rate-limiting enzyme of *de novo* steroidogenesis, we identify the constraint that maintains this equivalence. Cyp11a1 loss acts almost exclusively in females, where it simultaneously dampens the female-specific inflammatory programme and drives proliferation and IL-13 production above the level seen in either sex. Male Th2 cells are essentially unaffected. The effect requires differentiation, being absent in naive precursors, indicating that the steroidogenic machinery becomes functionally consequential only in the mature effector state. Cross-species analysis of two human atopic dermatitis single-cell datasets identifies a shared directional signature of gene expression. Cell-intrinsic steroidogenesis therefore constitutes a third axis of immunological sexual dimorphism, one that operates by uncoupling transcriptional identity from effector output rather than by specifying identity itself.

## INTRODUCTION

T helper 2 (Th2) cells are central orchestrators of type-2 immunity and key drivers of allergic diseases including asthma, atopic dermatitis, and food allergy^1–4^. Sexual dimorphism in type-2 immune responses is clinically significant. Women exhibit substantially higher prevalence and severity of allergic diseases compared with men^5–7^. Current mechanistic models attribute these sex differences primarily to two factors: circulating gonadal hormones and sex chromosome composition^8^. Both models, however, locate the source of dimorphism outside the responding cell: one in the endocrine environment, the other in chromosome dosage. Neither addresses whether a lymphocyte can generate sex-biased behaviour from a programme it runs itself. Whether immune cells possess such cell-autonomous mechanisms, and whether these act in the same direction as the systemic ones, remains unresolved.

We and others have previously demonstrated that Th2 cells express Cyp11a1 (also known as P450 side-chain cleavage enzyme or P450scc), the first and rate-limiting enzyme of the steroidogenesis pathway which converts cholesterol to pregnenolone, indicating that Th2 cells are capable of cell-intrinsic *de novo* steroid biosynthesis^9–13^. Cyp11a1 expression in T cells is induced during Th2 polarisation by type-2 cytokines including IL-4, and prior work has shown that cell-intrinsic steroidogenesis is functionally active in the differentiated Th2 state. The functional consequences of Cyp11a1 in T cells, however, have been examined without stratifying by sex, which leaves the direction of its contribution undetermined. Two opposite possibilities are consistent with existing data. If cell-intrinsic steroidogenesis sustains the female-biased Th2 state, its loss should reduce female Th2 effector output toward male levels. If instead it restrains that state, its loss should release female effector output beyond male levels. These predictions are readily distinguished, and they differ in what they imply for female-predominant allergic diseases^5–7^: the first casts steroidogenesis as a driver of pathology, the second as a protective brake whose failure would exacerbate it.

Here, we investigated whether T cell-intrinsic steroidogenesis contributes to sex-biased Th2 cell differentiation and function. Naïve CD4^+^ T cells from male and female T cell-specific *Cyp11a1* knockout and control mice were differentiated under identical *in vitro* Th2-polarising conditions and analysed using RNA sequencing, pathway-level analyses, functional assays, and cross-species comparison with human inflammatory-disease datasets. The second possibility proved correct, and with an unexpected asymmetry. Cyp11a1 loss shifted the female inflammatory transcriptional programme toward the male profile at the level of individual sex-dimorphic genes, while driving proliferation and IL-13 production above the level of either sex, dissociating transcriptional convergence from functional divergence within the same cells. Male Th2 cells were largely refractory. A subset of this signature overlapped directionally with female-biased transcriptional programmes in human atopic dermatitis.

## RESULTS

### Cyp11a1 loss remodels the female Th2 transcriptome, reducing inflammatory and increasing proliferative programmes

To define how cell-intrinsic steroidogenesis contributes to Th2 biology across sexes, we compared transcriptomes of control (*Cyp11a1*^fl/fl^) and Cyp11a1-deficient Th2 cells from female and male mice differentiated under identical *in vitro* condition (Figure 1A). Principal component analysis (PCA) revealed that control female Th2 cells were transcriptionally distant from control male Th2 cells along the first three principal components. *Cyp11a1* deletion markedly reduced the transcriptomic separation between female and male Th2 cells. Importantly, however, female knockout samples did not move into the male clusters: they occupied a distinct region of PCA space, indicating that loss of *Cyp11a1* releases a female-specific programme rather than converting female cells into male-like ones (Figure 1B). This shift indicates that Cyp11a1 acts on the female-biased transcriptional state, restraining its conversion into effector output in differentiated Th2 cells.

**Figure 1.**
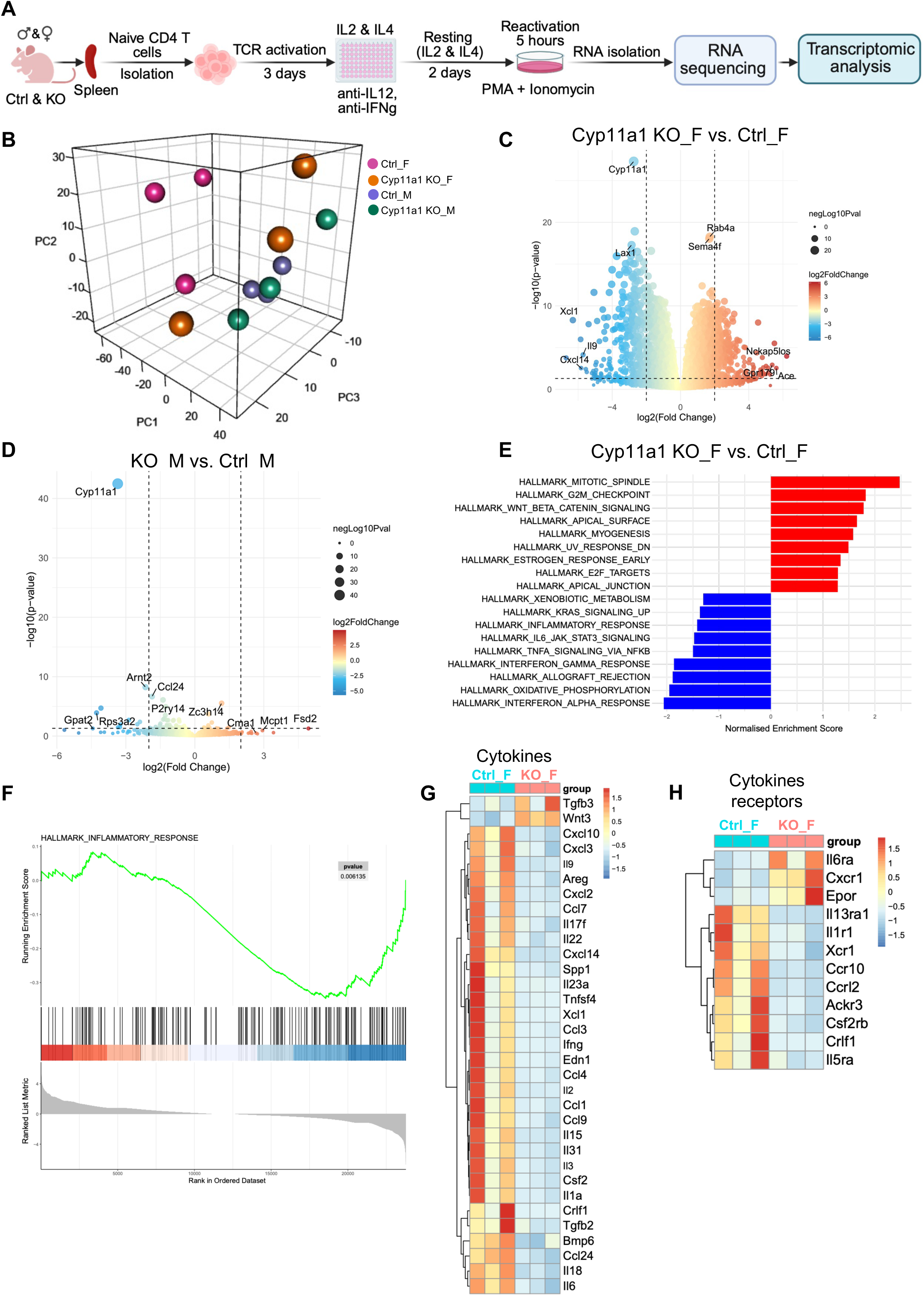
Cyp11a1 deletion preferentially reprogrammes the transcriptome of female Th2 cells. **(A)** Schematic representation of the experimental plan. Naïve CD4 T cells from *Cyp11a1*^fl/fl^ (ctrl) or *Cyp11a1* ^fl/fl^ *CD4*^Cre^ (KO) mice were isolated from spleen and activated for 3 days in anti-CD3e and CD28 antibody-coated plates in the presence of IL-2 and IL-4 with anti-IFNγ and anti-IL-12 antibodies. The cells were transferred to the resting plate for 2 days and reactivated with PMA and ionomycin for 5 hours. The cells were harvested, and RNA was prepared for sequencing. **(B)** Three-dimensional principal component analysis (PCA) of transcriptomes from *in vitro*-differentiated Th2 cells: Control female (Ctrl_F), Cyp11a1 KO female (Cyp11a1 KO_F), Control male (Ctrl_M), and Cyp11a1 KO male (Cyp11a1 KO_M). Note that Cyp11a1 KO female cells occupy a distinct region rather than converging on male clusters, consistent with loss of a female-specific constraint rather than acquisition of a male-like state. **(C)** Volcano plot showing differentially expressed genes (DEGs) in female Th2 cells (KO versus Ctrl). Significantly upregulated and downregulated genes are coloured by log₂ fold change; some of the top DEGs are labelled. **(D)** Volcano plot showing DEGs in male Th2 cells (KO versus Ctrl), demonstrating markedly fewer significant changes compared with females. **(E)** Hallmark pathway enrichment analysis in female Cyp11a1 KO versus Ctrl Th2 cells. Red bars indicate positively enriched pathways; blue bars indicate negatively enriched pathways. **(F)** GSEA enrichment plot for the HALLMARK_INFLAMMATORY_RESPONSE signature in female KO versus Ctrl comparison. **(G-H)** Heatmaps of differentially expressed cytokine (G) and cytokine receptor (H) genes in female Th2 cells (ctrl vs KO) with z-score-transformed expression values.

In females, *Cyp11a1* deletion produced broad gene-expression changes, as visualised by the female knockout-versus-control volcano plot, which showed numerous significantly altered transcripts, including depletion of *Cyp11a1* itself and marked changes in immune-related genes such as *Xcl1, Il9, Sema4a,* and *Rab4a* (Figure 1C, Supplementary Data 1). In contrast, the male knockout-versus-control volcano plot showed markedly fewer significant shifts, with *Cyp11a1* itself being the most prominently altered gene and only negligible additional changes in genes such as *Ccl24, Arnt2*, and a few immune-related transcripts (Figure 1D, Supplementary Data 2).

Pathway-level analysis showed this sex bias. In females, hallmark enrichment analysis indicated that *Cyp11a1* deletion dampened inflammatory programmes. These included signatures annotated as inflammatory response, IL-6/JAK–STAT3 signalling, TNFα signalling via NF-κB, IFN-α and -γ response, while concurrently enhancing cell-cycle-associated programmes, including G2M checkpoint, mitotic spindle, and Wnt/β-catenin signalling hallmarks (Figure 1E). Gene set enrichment analysis (GSEA) confirmed a significant downward shift in the “hallmark inflammatory response” signature in female *Cyp11a1* knockout versus control cells (Figure 1F). Differentially expressed cytokine and cytokine receptor genes in control female and *Cyp11a1* knockout female samples are visualised as a clustered heatmap (Figure 1G, 1H). Together, these data show that *Cyp11a1* deletion preferentially remodels the female Th2 transcriptome, with reduced inflammatory signatures and enhanced cell-cycle-associated programmes.

### Female and male Th2 cells differ in inflammatory programmes and in transcription factor activity at baseline

We next examined baseline sex differences in control (*Cyp11a1*^fl/fl^) female and male Th2 cells to establish the transcriptional framework upon which Cyp11a1 acts. The control female-versus-male volcano plot revealed robust sexually dimorphic transcription, including expected sex-linked transcripts such as high *Xist* signal on the female side and Y-linked genes (*Eif2s3y, Ddx3y, Kdm5d, Uty*) on the male side (Figure 2A, Supplementary Data 3). Numerous autosomal immune-associated genes, including *Fcer1a* and *Slc25a31*, were found differentially expressed (Figure 2A). Hallmark analysis further demonstrated that sex differences in control Th2 extended well beyond sex-chromosome genes. Female Th2 cells were enriched for IFNα and γ response, IL-6/JAK–STAT3 signalling, allograft rejection, and TNFα signalling via NF-κB pathways, while male cells showed enrichment for epithelial-mesenchymal transition, G2M checkpoint, myogenesis, and mitotic spindle programmes (Figure 2B). This pattern indicates that female Th2 cells display heightened immune activation signatures relative to male cells under identical differentiation conditions.

**Figure 2.**
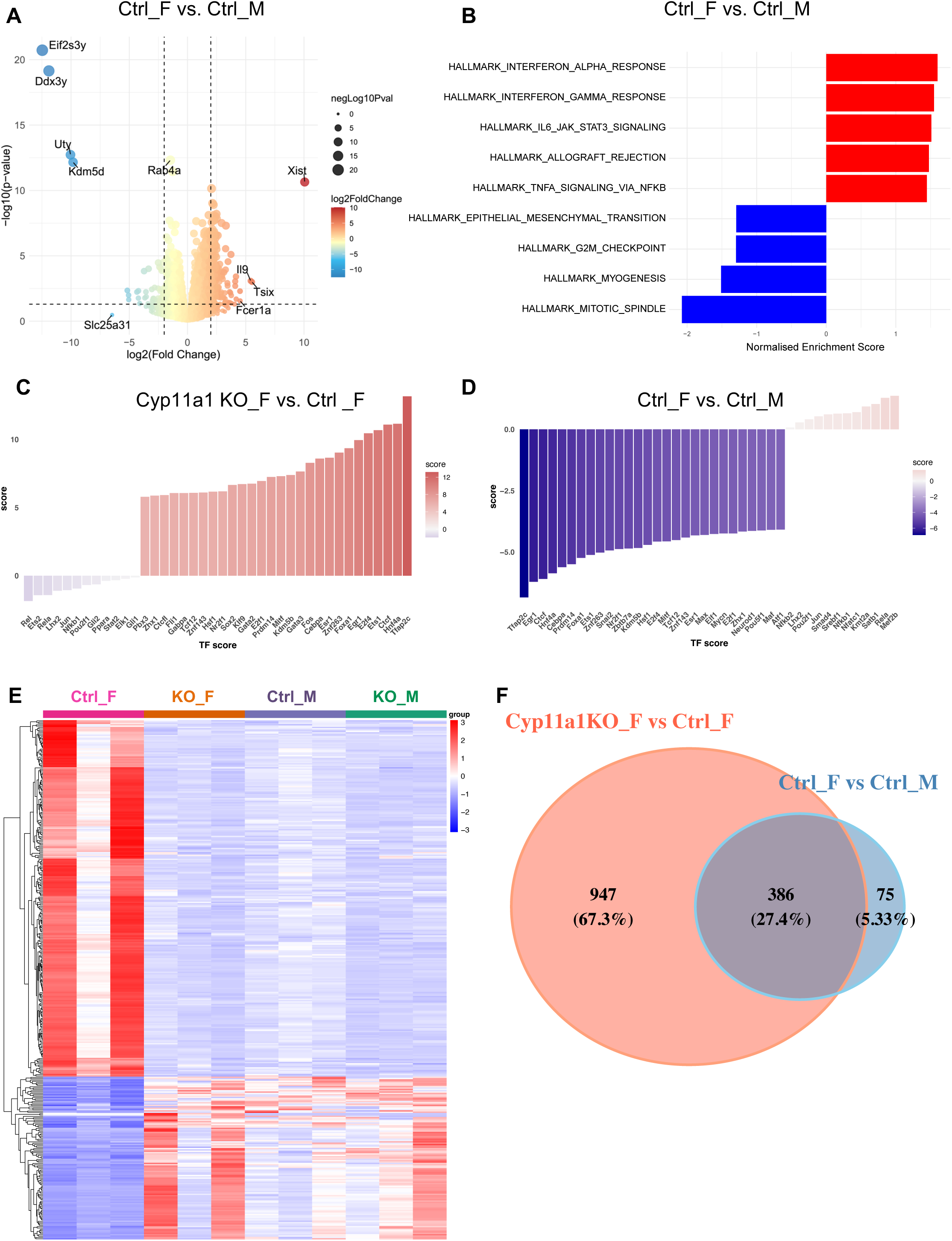
Cell-intrinsic steroidogenesis programmes female Th2 identity. **(A)** Volcano plot showing differentially expressed genes (DEGs) between control female (Ctrl_F) and control male (Ctrl_M) Th2 cells. **(B)** Hallmark pathway enrichment analysis comparing control female (Ctrl_F) versus control male (Ctrl_M) Th2 cells. Red bars: pathways enriched in females; blue bars: pathways enriched in males. **(C)** Transcription factor (TF) activity scores for genes differentially regulated in Cyp11a1 knockout female Th2 compared to control female Th2. **(D)** Transcription factor (TF) activity scores for genes differentially regulated in control female versus control male Th2 cells. **(E)** Hierarchical clustering heatmap of all four groups (Ctrl_F, Cyp11a1 KO_F, Ctrl_M, Cyp11a1 KO_M) based on sex-dimorphic and Cyp11a1-dependent DEGs, displayed as z-score-normalised expression values. **(F)** Venn diagram showing overlap between DEGs from the female KO versus control comparison (Cyp11a1KO_F vs Ctrl_F) and the control female versus control male comparison (Ctrl_F vs Ctrl_M). Of the control sex-dimorphic signature (461 genes), 386 overlaps with *Cyp11a1*-dependent genes.

To interrogate the regulatory architecture underlying these differences, we performed transcription factor (TF) activity scoring on two separate contrasts. In female Th2 cells, *Cyp11a1* deletion shifted the inferred activity of a distinct set of TFs relative to female controls, with the largest changes among NF-κB family members (*Rel, Rela, Nfkb1*) and AP-1 components (Figure 2C). Independently, comparison of control female with control male Th2 cells showed that an equally broad TF panel already distinguishes the two sexes before any genetic perturbation (Figure 2D). The overlap between the two panels indicates that Cyp11a1 acts on the same regulatory nodes that establish baseline sex dimorphism, rather than on a separate downstream layer.

Hierarchical clustering of all four groups (control female, *Cyp11a1* knockout female, control male, *Cyp11a1* knockout male) using both sex-dimorphic and Cyp11a1-dependent differentially expressed genes (DEGs) demonstrated that within this sex-dimorphic gene set specifically, *Cyp11a1* deletion shifted female Th2 cells toward male expression levels (Figure 2E, Supplementary Data 4). Venn diagram analysis quantified this convergence. Out of 461 genes constituting the control female-versus-male sex-dimorphic signature, 386 were also significantly altered by *Cyp11a1* deletion in females, with only 75 genes remaining uniquely sex-dimorphic independently of steroidogenesis (Figure 2F). The Cyp11a1-dependent gene set included an additional 947 genes not part of the baseline sex signature, indicating that steroidogenic disruption also perturbs programmes beyond the canonical sex-dimorphic transcriptome. Because gene-set overlap alone cannot distinguish a directed shift from a nonspecific perturbation, we assessed overlap together with the directionality of the hierarchical clustering in Figure 2E. The two analyses agreed: Cyp11a1-dependent changes in females occur predominantly in sex-dimorphic genes and move them toward the male expression level. Cyp11a1-dependent steroidogenesis is therefore a major determinant of the female-biased Th2 transcriptional programme, though as shown below this transcriptional convergence does not extend to effector function.

### Steroidogenic regulation requires Th2 differentiation and is absent in naive precursors

To determine whether the sex-biased impact of Cyp11a1 is already present in naïve precursors or instead emerges during Th2 differentiation, we analysed naïve CD4+ T cell transcriptomes from the same four experimental groups (Figure 3A). PCA of naïve T cell samples indicated that sex was the dominant axis of variance along PC1, whereas genotype effects appeared comparatively subtle, with control (*Cyp11a1*^fl/fl^) and *Cyp11a1*-knockout samples of the same sex remaining in close proximity in PCA space (Figure 3B). In naive cells, *Cyp11a1* deletion had little transcriptional consequence in either sex. The female knockout-versus-control comparison yielded few significantly altered genes, a markedly smaller landscape than the corresponding Th2 comparison (Figure 3C, Supplementary Data 5), and the male comparison was similarly sparse (Figure 3D, Supplementary Data 6). By contrast, the control female-versus-male naive comparison was dominated by sex-chromosome-linked transcripts (*Xist, Eif2s3y, Ddx3y, Kdm5d, Uty*) with limited autosomal immune divergence (Figure 3E, Supplementary Data 7), confirming that naive cells of the two sexes differ chiefly in sex-linked gene dosage rather than in immune programme. Cyp11a1-dependent regulation is therefore not pre-established in precursors but emerges with Th2 differentiation.

**Figure 3.**
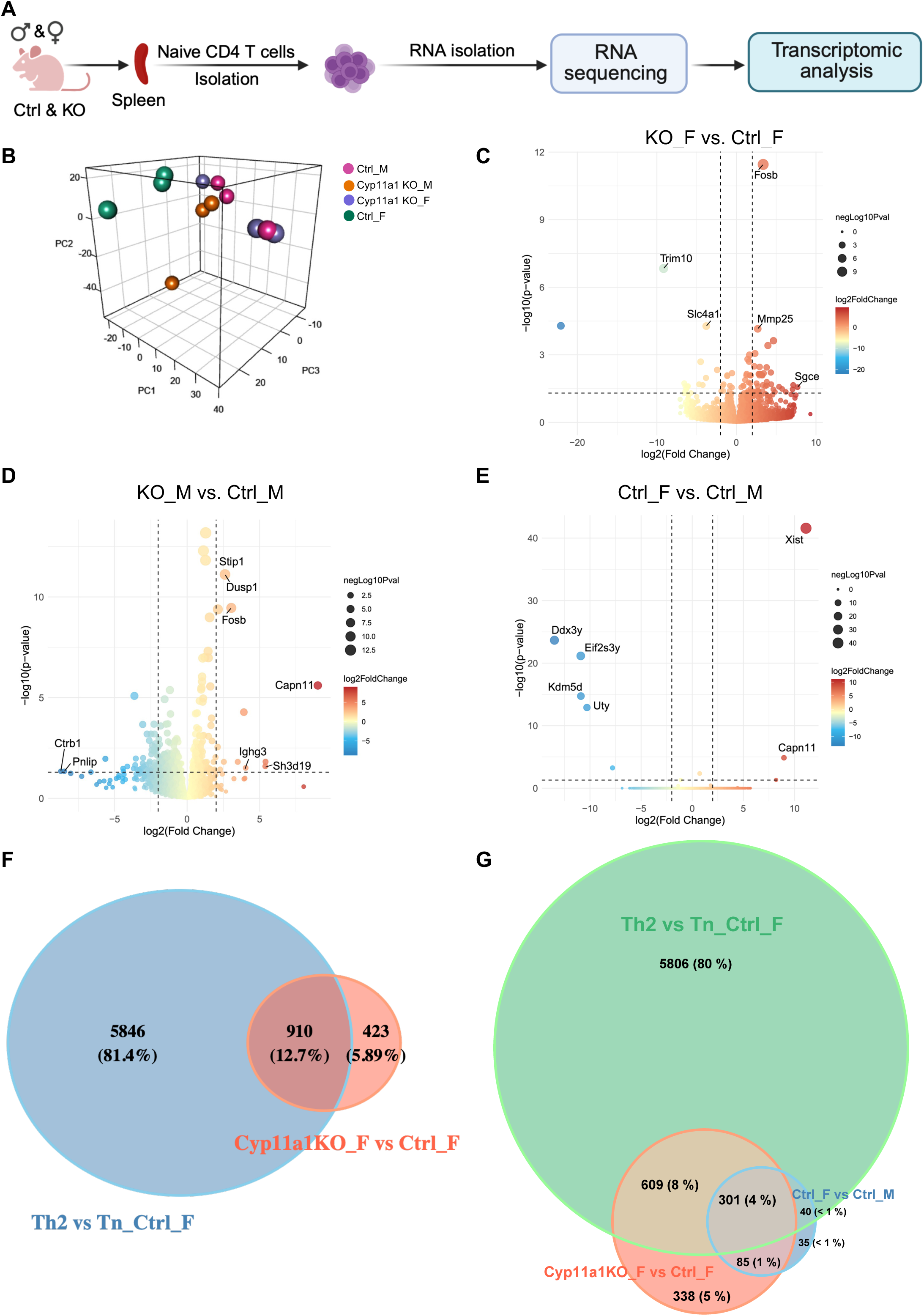
Steroidogenic regulation is developmentally stage-specific. **(A)** Schematic diagram of the experimental plan for naïve T cell transcriptomic analysis. **(B)** Three-dimensional PCA of naïve T cell transcriptomes from all four groups: *Cyp11a1*^fl/fl^ (ctrl) male (M); Cyp11a1 KO male; *Cyp11a1*^fl/fl^ (ctrl) female (F); Cyp11a1 KO female. Sex is the dominant axis of variance; genotype effects are comparatively subtle. **(C)** Volcano plot of naïve female Cyp11a1 KO versus control naïve CD4+ T cells, showing few significantly altered genes, the most prominent being Fosb, Trim10, Slc4a1, Mmp25 and Sgce. **(D)** Volcano plot of naïve male Cyp11a1 KO versus control naive CD4+ T cells. **(E)** Volcano plot of naïve control female versus control male CD4+ T cells. **(F)** Venn diagram comparing Th2 differentiation-associated DEGs (Th2 versus naïve in control females) with Cyp11a1-dependent DEGs (KO versus control in female Th2 cells), showing that Cyp11a1-dependent genes represent a specific subset of the differentiation programme. **(G)** Three-way Venn diagram showing the relationship between differentiation-associated DEGs (Th2 versus naïve in Ctrl_F), Cyp11a1-dependent DEGs (KO versus control in female Th2 cells), and sex-dimorphic DEGs (Ctrl_F versus Ctrl_M).

Overlap analyses provided quantitative support for stage-specific regulation. Comparing the differentiation-associated transcriptional programme (Th2 versus naïve in control females; 6,756 DEGs) with the Cyp11a1-dependent programme (knockout versus control in female Th2; 1,333 DEGs) revealed that 910 genes were shared, while most differentiation genes (5,846) were Cyp11a1-independent and 423 genes were uniquely Cyp11a1-dependent (Figure 3F). A three-way comparison incorporating the control sex-dimorphic gene set revealed the specificity of this relationship: of the 461 genes distinguishing control female from control male Th2 cells, 386 (83.7%) were also altered by *Cyp11a1* deletion in females (Figure 3G). Cyp11a1-dependent genes therefore do not merely overlap the sex-dimorphic programme but encompass the large majority of it, indicating that *de novo* steroidogenesis acts on the sex-dimorphic transcriptional programme itself rather than in parallel with it. . These findings support a model in which Cyp11a1-dependent steroidogenesis acts primarily during or after Th2 polarisation to shape mature-state programmes rather than broadly reprogramming naïve precursors.

### Cyp11a1 loss selectively enhances proliferation and IL-13 production in female Th2 cells

Transcriptomic analysis had revealed coordinate dysregulation of genes governing G2M checkpoint progression, mitotic spindle assembly, and proliferative control, suggesting that steroidogenic disruption would alter Th2 cell expansion kinetics. To test this prediction, naïve CD4^+^ T cells were labelled with CellTrace Violet (CTV) prior to Th2 activation and differentiation, and proliferation was quantified by flow cytometric analysis of dye decay (Figure 4A). Female *Cyp11a1*-deficient Th2 cells exhibited significantly enhanced proliferative capacity compared with controls (Figure 4B). In contrast, male *Cyp11a1*-knockout Th2 cells showed no significant change in division index (Figure 4C). Importantly, the male refractoriness cannot be attributed to inefficient deletion: *Cyp11a1* transcript was reduced to a comparable degree in male and female knockout Th2 cells (Figure 1D), establishing that the gene was effectively ablated in both sexes and that the divergent functional outcome reflects a genuine difference in downstream dependence rather than differential recombination efficiency.

**Figure 4.**
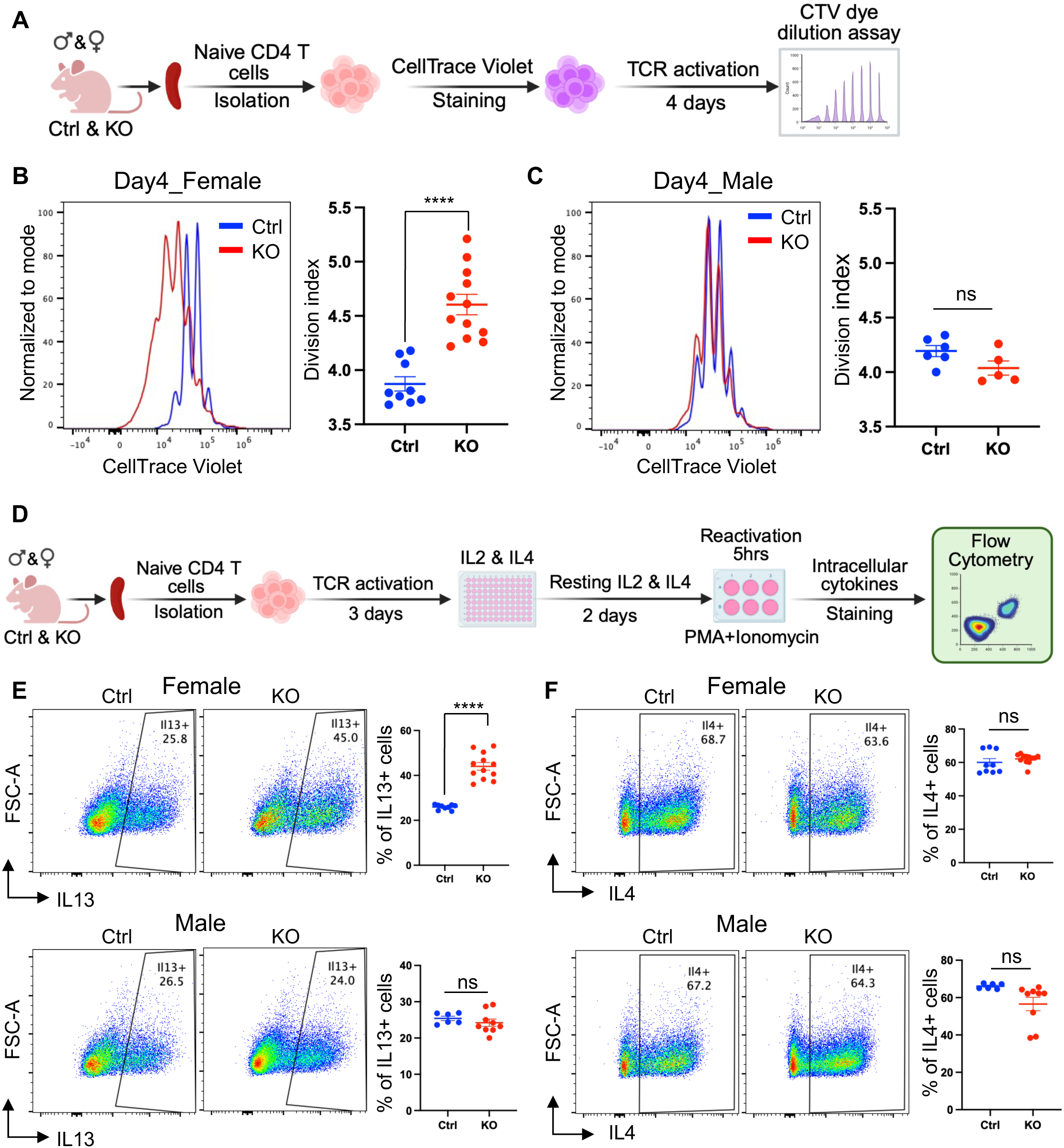
Cyp11a1 loss selectively enhances proliferation and IL-13 production in female Th2 cells. **(A)** Schematic of the CellTrace Violet (CTV) proliferation assay. Naïve CD4⁺ T cells from control and Cyp11a1 KO mice were labelled with CTV and differentiated under Th2-polarising conditions for 4 days, after which CTV dye dilution was measured by flow cytometry. **(B)** CTV proliferation assay in female Th2 cells. Left: representative CTV decay histograms for control (blue) and Cyp11a1 KO (red) female Th2 cells at day 4. Right: quantification of division index. Female KO Th2 cells show significantly enhanced proliferation (****p < 0.0001). Each dot represents one independent biological sample (n=9-13). **(C)** CTV proliferation assay in male Th2 cells. Left: representative histograms. Right: division index quantification showing no significant change in male KO Th2 cells compared with controls (ns, p > 0.05). Each dot represents one independent biological sample (n=5-6). **(D)** Schematic of the intracellular cytokine staining assay. Following Th2 differentiation, cells were reactivated with PMA and ionomycin for 5 hours (monensin added last 3 hours to block protein secretion) and stained for intracellular IL-13 and IL- 4. **(E)** Left: representative flow cytometry dot plots for female (top) and male (bottom) control and KO Th2 cells. Right: quantification of % IL-13^+^ Th2 cells. Female KO cells exhibit significantly elevated IL-13 production (****p < 0.0001); male KO cells show no significant change (ns). Each dot represents one independent biological sample (n=6-11). **(F)** Left: representative flow cytometry dot plots of IL-4 expression in female (top) and male (bottom) control and Cyp11a1 KO Th2 cells. Right: quantification of % IL-4⁺ cells. Female and male KO cells do not show significant change in IL-4 expression (ns, p >0.05). Each dot represents one independent biological sample (n=6-11). All error bars indicate mean ± SEM. P values were calculated using two-tailed unpaired Student’s t test.

In T helper cell differentiation, proliferation is intrinsically coupled with lineage commitment and effector function acquisition^14–16^. The number of cell divisions often correlates with cytokine production capacity, as successive divisions drive progressive chromatin remodelling at cytokine loci^14^. Therefore, we next tested whether the enhanced proliferation observed in female Cyp11a1-deficient cells was accompanied by altered cytokine expression (Figure 4D). Intracellular cytokine staining confirmed this prediction. Female *Cyp11a1* knockout Th2 cells expressed significantly elevated levels of IL-13 compared to control female Th2 (Figure 4E, top panel). Male knockout cells showed no significant change in IL-13 expression (Figure 4E, bottom panel). Notably, this cytokine-specific dysregulation did not extend equally to all Th2 effector cytokines. IL-4 expression showed no significant increase in female knockout cells (Figure 4F, top panel) or in male knockout cells (Figure 4F, bottom panel).

### Female Cyp11a1-deficient mouse Th2 signatures partially overlap with female-biased human T helper states in type-2 inflammatory skin disease

To assess disease relevance, female murine Cyp11a1-dependent Th2 signatures (Supplementary Data 1) were compared with sex-stratified transcriptomes of CD4^+^CD3E^+^GATA3^+^ T helper cells from atopic dermatitis (GSE204762)^17^ and (GSE213849)^18^ scRNA-seq datasets (Figure 5A, B), and female-versus-male differential expression was analysed separately for each disease dataset (Figure 5C, D). This analysis was designed to test whether genes increased after *Cyp11a1* deletion in female mouse Th2 cells show directional overlap with genes increased in female human T helper cells in type-2 inflammatory disease.

**Figure 5.**
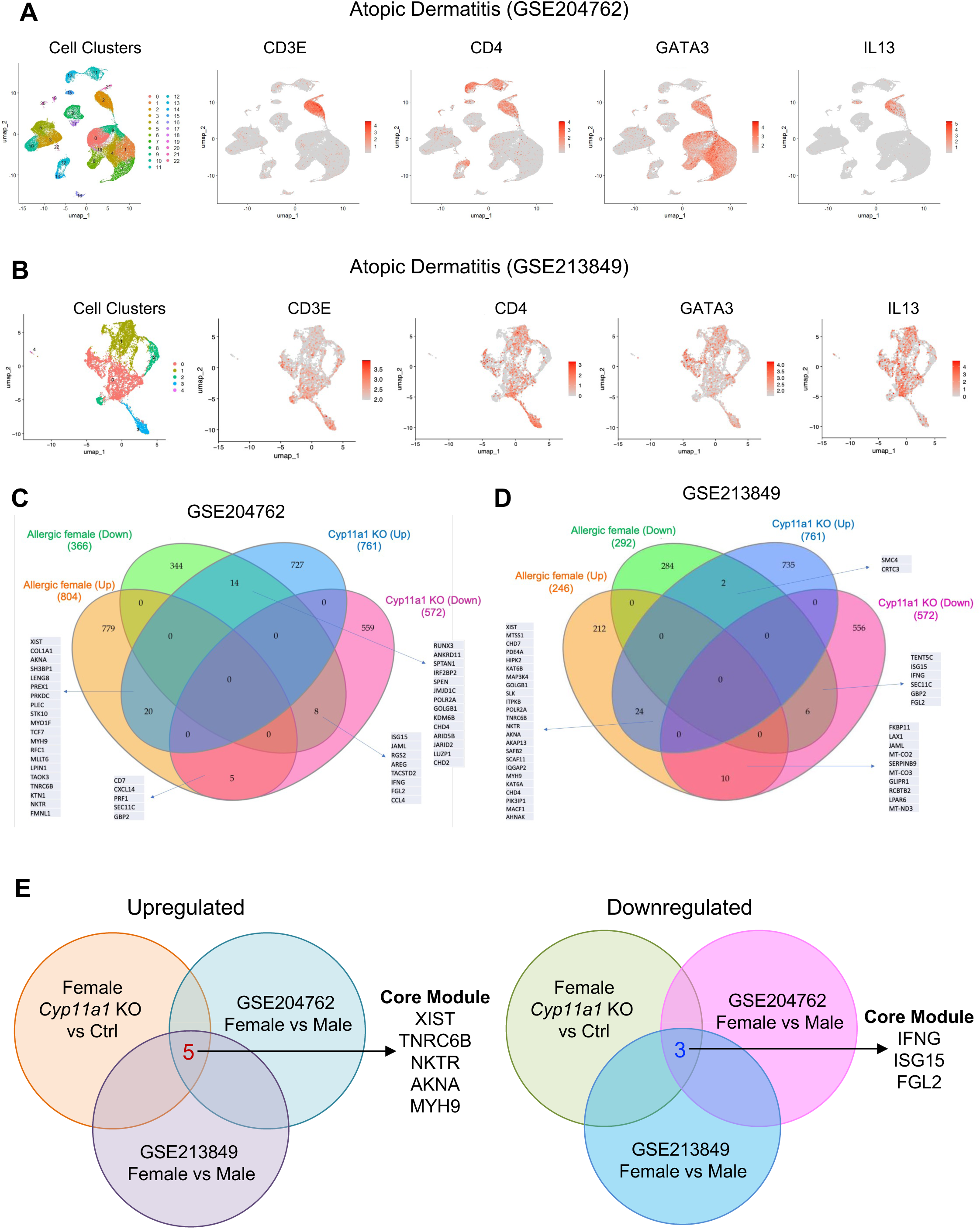
A subset of the female *Cyp11a1*-dependent murine Th2 transcriptional signature overlaps directionally with female-biased human Th2 gene expression in type-2 inflammatory skin disease (atopic dermatitis). **(A-B)** UMAP embedding of single-cell transcriptomes from human atopic dermatitis skin dataset (GSE204762) (A) and (GSE213849) (B), showing the major cell clusters identified after Seurat-based integration and clustering. The *CD4^+^CD3E^+^GATA3^+^*T helper cell population was selected for downstream sex-stratified differential expression analysis. *IL13* expression was verified to confirm their allergic Th2 signature. **(C-D)** Four-way Venn diagrammes comparing differentially expressed gene sets from female *Cyp11a1* KO versus control mouse Th2 cells (upregulated, blue: 761 genes; downregulated, magenta: 572 genes) with DEGs from female versus male *CD4*^+^*CD3E^+^GATA3^+^*T helper cells in human atopic dermatitis (GSE204762; upregulated, orange: 804 genes; downregulated, green: 366 genes) (C) and (GSE213849; upregulated, orange: 246 genes; downregulated, green: 292 genes) (D). Numbers indicate the size of each intersection, and selected genes from representative overlap regions are shown. **(E)** Left panel: five genes were common to the mouse *Cyp11a1* KO-upregulated female Th2 set and the female-upregulated human T helper sets in both datasets: XIST, *TNRC6B, NKTR, AKNA* and *MYH9*. Of these, *XIST* serves as an internal sex-assignment control, confirming correct sex annotation in the human datasets; the four remaining genes (*TNRC6B, NKTR, AKNA, MYH9*) constitute the conserved upregulated module. Right panel: the three genes common to the mouse *Cyp11a1* KO-downregulated female Th2 set and the female-downregulated human T helper sets in both datasets were *IFNG, ISG15*, and *FGL2*. Together, these gene modules define the conserved concordant transcriptional overlap between female murine *Cyp11a1* deficiency and female-biased human T helper states across both type-2 inflammatory skin disease datasets analysed. Note that the murine comparison is a genotype contrast within females (Cyp11a1 KO versus control), whereas both human comparisons are female-versus-male contrasts within disease; the overlap therefore tests whether genes de-repressed by loss of steroidogenic restraint in mouse correspond to genes elevated in female human T helper cells.

In atopic dermatitis, 804 genes were upregulated and 366 genes were downregulated in female compared with male Th2 cells (Figure 5C). Here, the human female-upregulated set showed a 20-gene overlap with the mouse *Cyp11a1* KO-upregulated set, along with additional smaller intersections involving the downregulated sets (Figure 5C). In another dataset of atopic dermatitis, 246 genes were upregulated and 292 genes were downregulated in female compared with male Th2 cells (Figure 5D). Comparison of these gene sets with the female mouse *Cyp11a1* knockout Th2 signature showed a 24-gene overlap between the human female-upregulated set and the mouse *Cyp11a1* KO-upregulated set, together with additional smaller overlaps involving the corresponding downregulated sets (Figure 5D).

Intersecting same-direction overlaps across both human datasets identified five shared upregulated genes: *TNRC6B, NKTR, AKNA, MYH9* and *XIST* (Figure 5E, left panel). Because *XIST* is necessarily elevated in female relative to male human cells irrespective of disease state, we treat it as an internal control for sex assignment rather than as an informative component of the module, and consider the conserved signal to comprise the four remaining genes. These four represent the shared transcriptional signal linking female murine Cyp11a1-deficient Th2 cells with female-biased human T helper states across the two disease (atopic dermatitis) datasets examined. The functional identities of the four conserved genes are collectively consistent with a state of heightened female-biased T helper activation. We note in passing that *XIST*, although used here as a sex-assignment control, is itself known to form ribonucleoprotein complexes that promote sex-specific epigenetic immune activation in T cells^19, 20^; its recovery in both datasets is therefore reassuring but not independent evidence for the module. *TNRC6B* encodes the GW182 scaffold protein of the miRNA-induced silencing complex and its upregulation implies broad post-transcriptional rewiring, potentially stabilising pro-inflammatory transcripts and suppressing negative immune regulators^21^. *AKNA* co-ordinately regulates CD40 and CD40 ligand, thereby promoting T–B cell collaboration and IgE class switching, a process directly relevant to elevated IgE and chronic Th2 responses characteristic of atopic dermatitis. *NKTR* is implicated in IL-2 pathway amplification and effector T cell survival, while *MYH9*, encoding non-muscle myosin IIA, supports immune synapse maturation and T cell receptor-coupled calcium signalling. Taken together, this four-gene module (considered alongside XIST as a sex-assignment control) is consistent with a female Th2-associated programme characterised by sex-linked epigenetic regulation, post-transcriptional control of inflammatory gene expression, enhanced co-stimulatory potential, and cytoskeletal features associated with T cell activation. Its shared upregulation across murine *Cyp11a1*-deficient Th2 cells and female human GATA3^+^ T helper cells in two independent type-2 skin disease datasets suggests that these functional attributes may form part of a conserved transcriptional correlate of reduced steroidogenic restraint.

Cross-species integration also identified a conserved three-gene module, *ISG15, IFNG*, and *FGL2*, that was downregulated (Figure 5E, right panel). Functionally, these genes are associated with interferon-responsive and immunoregulatory programmes: IFNG encodes the canonical Th1 cytokine that antagonises Th2 responses, ISG15 is an interferon-stimulated type-1 immune regulatory gene, and FGL2 has been linked to suppressive immune regulation including Treg effector function. Their shared reduction across all three conditions is therefore consistent with attenuation of counter-regulatory pathways that would normally constrain type 2 inflammation.

## DISCUSSION

This study identifies Cyp11a1-dependent steroidogenesis as a cell-intrinsic checkpoint that uncouples the transcriptional identity of female Th2 cells from their effector output. The finding rests on an observation: female and male Th2 cells differentiated under identical hormone-free conditions are transcriptionally distinct, with female cells carrying a heightened inflammatory and interferon-response programme, yet they produce IL-13 and IL-4 at indistinguishable rates, differing only modestly in baseline proliferative capacity (Supplementary Figure S1). A difference in programme without a difference in output implies that something converts the former into the latter at a controlled rate, and that this control is sex-specific. Cyp11a1 is that control. Its deletion in females simultaneously dampened the inflammatory programme and drove proliferation and IL-13 production above the level of either sex, whereas the same deletion in males was almost without effect. This is what would be expected if the pathway constrains a programme that males do not possess. Cyp11a1 loss therefore does not masculinise female Th2 cells. It removes a brake, and the resulting state is not male-like but hyper-functional relative to both sexes. The male phenotype is therefore best understood as an absence of substrate for the mechanism rather than an absence of the mechanism itself. *Cyp11a1* was deleted with comparable efficiency in both sexes (Figure 1D), yet only female cells responded, indicating that the steroidogenic checkpoint is engaged only where a female-biased inflammatory programme exists to be restrained. Because this occurs in cells differentiated in the absence of circulating gonadal hormones, the mechanism is cell-autonomous and separable from the two axes to which immunological sexual dimorphism is conventionally attributed.

Differentiating naive CD4⁺ T cells under identical *in vitro* conditions removes circulating gonadal hormones as a variable, and under these conditions a robust transcriptional sex dimorphism persists — establishing that a substantial component of Th2 sexual dimorphism is carried by the cell rather than supplied by its endocrine environment. Genes distinguishing female from male Th2 cells were preferentially altered by *Cyp11a1* deletion in females, indicating that *de novo* steroidogenesis acts on the sex-dimorphic programme itself rather than alongside it. We note that this design removes only acute hormonal exposure; it does not exclude durable effects of prior *in vivo* hormone exposure imprinted on the precursor pool before isolation. These observations extend prior work showing that Cyp11a1 expression is induced during IL-4-driven polarisation and are consistent with a model in which the steroidogenic machinery becomes functionally embedded in the differentiating Th2 cell state. We and others have previously shown that immune cells (Th2, Th17, macrophages and mast cells) synthesise steroid pregnenolone in a Cyp11a1-dependent manner, establishing that loss of this enzyme eliminates the pathway’s biochemical output^9–13, 22–24^.

Current models of immunological sexual dimorphism invoke two principal mechanisms: gonadal hormones and sex chromosome complement^25^. Oestrogen enhances Th2 responses^25^, while androgens suppress type-2 inflammation^26^. In addition, the four-core genotype model has shown X-chromosome dose effects, particularly through genes that escape X-inactivation such as TLR7 ^27, 28^. Our findings introduce a third mechanism that is located in neither the endocrine environment nor chromosome dosage: *de novo* steroid synthesis within the immune cell itself. This axis is distinguished from the other two not only by its location but by its logic. Gonadal hormones and X-linked gene dosage are generally described as amplifying female-biased type-2 responses; the steroidogenic axis described here restrains them, and does so specifically in females. That the two classes of mechanism act in opposing directions suggests they are not redundant, and that female type-2 immunity reflects a balance between them rather than the sum of parallel drivers.

One possible interpretation is that loss of Cyp11a1-dependent steroidogenesis releases a proliferative restraint in female Th2 cells, and that additional divisions contribute to enhanced IL-13 competence through progressive chromatin remodelling at cytokine loci. However, the present experiments do not directly distinguish whether altered division history drives increased IL-13 output or whether cytokine dysregulation occurs through a partly independent transcriptional mechanism. The selective elevation of IL-13 over IL-4 is notable, because IL-13 is a dominant effector cytokine in airway remodelling, mucus hypersecretion, and tissue fibrosis in asthma and atopic disease. This raises the possibility that sex-differential dysregulation of Th2-intrinsic steroidogenesis could be relevant to variation in disease expression or therapeutic responsiveness, although that question remains to be tested directly.

The stage-specificity of steroidogenic regulation, with minimal transcriptional effects in naïve T cells but extensive reprogramming in differentiated Th2 cells, demonstrates that this pathway engages during or after polarisation rather than being pre-established in precursors. This is consistent with prior evidence that Cyp11a1 expression is induced under IL-4-polarising conditions^9, 10^, suggesting that the steroidogenic machinery is assembled during the differentiation programme and becomes functionally important specifically in the mature Th2 effector state.

Cross-species comparison identified a four-gene module (i.e., *TNRC6B, NKTR, AKNA*, and *MYH9*, identified alongside *XIST* as an internal control for sex assignment) that was concurrently upregulated in female Cyp11a1-deficient mouse Th2 cells and in female Th2 cells from two independent human type-2 inflammatory skin disease datasets. Collectively, these genes are compatible with enhanced sex-linked epigenetic activity, post-transcriptional rewiring of pro-inflammatory transcripts, co-stimulatory T-B cell signalling, effector T cell survival, and cytoskeletal features of T cell activation. Reciprocally, *IFNG, ISG15*, and *FGL2* were consistently downregulated across the murine and human comparisons, consistent with attenuation of counter-regulatory programmes that would normally constrain type-2 inflammation. The human analyses are correlative and do not demonstrate impaired CYP11A1 activity or causal steroidogenic dependence in patient T helper cells. The shared gene modules are therefore best viewed as plausible candidate readouts for future mechanistic studies rather than evidence of an established conserved pathway in human disease.

These findings have potential translational relevance for female-predominant allergic and type-2 inflammatory diseases. Cell-intrinsic steroidogenesis represents a regulatory axis conceptually distinct from systemic gonadal hormone exposure, and one that could in principle be targeted independently of systemic endocrine manipulation. Future studies should test whether restoring Cyp11a1 activity, or modulating its steroidogenic products, can alter pathogenic Th2 proliferation and IL-13 production in preclinical disease models. CYP11A1 expression or activity in T helper cells may also warrant evaluation as a candidate biomarker in sex-stratified clinical settings, although prospective validation will be required before such applications can be considered.

Two limitations bound the interpretation of this work. First, all murine data derive from in vitro-differentiated Th2 cells; we have not tested whether female Cyp11a1 deficiency alters the course of type-2 inflammation *in vivo*, and the functional consequences we observe may be modified by the tissue environment, by antigen persistence, or by regulatory populations absent from these cultures. Second, the human analyses are cross-sectional comparisons between sexes in disease tissue and cannot establish that CYP11A1 activity is altered in patient T helper cells; the shared modules are candidate readouts for future mechanistic study rather than evidence of a validated conserved pathway. Each of these defines a specific experiment, and none undermines the central *in vitro* genetic result: that Cyp11a1 constrains effector output in female but not male Th2 cells.

In summary, cell-intrinsic steroidogenesis emerges as a sex-specific determinant of Th2 effector identity that operates independently of circulating hormones. These findings argue for a reconceptualisation of immunological sexual dimorphism that incorporates cell-autonomous steroid metabolic programmes alongside established gonadal and chromosomal mechanisms and identify Cyp11a1 as a candidate target for sex-stratified therapeutic intervention in type-2 allergic disease.

## METHODS

### Mice

All mice were maintained and used in accordance with the UK Animals (Scientific Procedures) Act 1986 Amendment Regulations 2012 and the UK Animals in Science Regulation Unit’s Code of Practice for the Housing and Care of Animals Bred, Supplied, or Used for Scientific Purposes. All experimental protocols were conducted under a UK Home Office Project Licence (PPL PP4938782) and received approval from the University of Cambridge Animal Welfare and Ethical Review Body. Sample sizes were determined based on prior experience and a priori power analysis (G*Power). Mice were housed in a specific pathogen-free facility at the Gurdon Institute on a 12-hour light/dark cycle and genotyped by Transnetyx.

*Cyp11a1*^fl/fl^ mouse line was generated at the Wellcome Sanger Institute as previously described^10^. T cell specific knockout mice (*Cyp11a1*^fl/fl^;*Cd4*^Cre^) were generated by crossing *Cyp11a1*^fl/fl^ with *Cd4*^Cre^ mice (Jackson laboratory). In this study we used mice aged between 8 to 12 weeks. Male and female mice were age-matched within experiments and analysed as independent biological replicates.

### Naïve T cell isolation and Th2 differentiation

Mouse spleens were harvested and mechanically dissociated through a 70-μm nylon cell strainer to obtain single-cell suspensions. Red blood cells were lysed using 1× RBC lysis buffer (eBioscience) for 3 minutes at room temperature, quenched with PBS, and washed. Naïve CD4+CD62L+ T cells were isolated using the Naïve CD4+ T Cell Isolation Kit II (Miltenyi Biotec) according to the manufacturer’s instructions.

Purified naïve T cells were activated on plates pre-coated with anti-CD3e (1 μg/mL, clone 145-2C11, eBioscience) and anti-CD28 (5 μg/mL, clone 37.51, eBioscience) under Th2-polarising conditions: recombinant murine IL-4 and IL-2 (10 ng/mL, R&D Systems), anti-IL-12 (5 μg/mL, eBioscience) and anti-IFN-γ (5 μg/mL, clone XMG1.2, eBioscience). Cells were cultured at 37°C in a humidified incubator with 5% CO₂. After 72 hours, cells were transferred to resting plates with fresh medium containing IL-2 and IL-4 for an additional 48 hours. For cytokine analysis, cells were reactivated with PMA and ionomycin for 5 hours; monensin was added after the first two hours for the remaining 3 hours. Cells were then harvested for intracellular cytokine staining.

### Spectral flow cytometry

Surface and intracellular cytokine staining were performed according to the manufacturer’s instructions using the eBioscience intracellular staining workflow. Briefly, single-cell suspensions were stained with Live/Dead Fixable Dead cell stain kit (Molecular Probes/ Thermo Fisher) and blocked by purified rat anti-mouse CD16/CD32 purchased from BD Bioscience and eBioscience. Surface staining was performed in flow cytometry staining buffer (eBioscience) or in PBS containing 3% FCS at 4 °C followed by intracellular cytoplasmic protein staining overnight at 4 °C. After staining, cells were washed with flow cytometry staining buffer (eBioscience) or 3% PBS-FCS, and were analysed by Cytek Aurora (5L) flow cytometer. FlowJo v10.10.0 software facilitated the data analysis.

### Proliferation assay

Naïve CD4+ T cells from control and Cyp11a1 knockout mice were labelled with Cell Trace Violet (CTV) according to the manufacturer’s protocol (Invitrogen) before activation under Th2-polarising conditions as described above. After 4 days of culture, CTV fluorescence decay was assessed by spectral flow cytometry on a Cytek Aurora instrument. Division index was calculated using the proliferation modelling tools in FlowJo.

### RNA sequencing

#### Experimental design

Two distinct T cell populations were analysed: naïve T cells and *in vitro*-differentiated Th2 cells. Naïve CD4+CD62L+ T cells were sorted from spleens of age-matched mice (8–12 weeks) into four experimental groups: female control (Ctrl_F), female Cyp11a1 knockout (Cyp11a1KO_F), male control (Ctrl_M), and male Cyp11a1 knockout (Cyp11a1KO_M). Each group contained three independent biological replicates (n = 3 per group).

#### RNA extraction and library preparation

Total RNA was extracted using the RNeasy Plus Mini Kit (Qiagen). RNA quality was assessed by NanoDrop spectrophotometry (A260/A280 and A260/A230 ratios). mRNA was purified using poly-T oligo-attached magnetic beads, fragmented, and converted to cDNA using random hexamer primers. Libraries were constructed by end repair, A-tailing, adapter ligation, size selection, and PCR amplification, then validated by Qubit, real-time PCR, and Bioanalyzer.

#### Sequencing and alignment

Sequencing was performed on a NextSeq 500 platform (Illumina). Raw reads were quality-controlled using FastQC and aligned to the mouse reference genome GRCm38.p4 (Gencode M10) using HISAT2 (v2.2.1). SAM files were converted and sorted to BAM format using SAMtools. Gene-level counts were generated using htseq-count.

#### Differential expression analysis

Differential gene expression was analysed using DESeq2 (v1.34.0) within R (v4.2.1) ^29^. A DESeqDataSet object was created with a design formula (∼group) incorporating the four experimental conditions. Size factors were estimated using the estimateSizeFactors function, followed by variance stabilisation and dispersion estimation with default parameters. Pairwise contrasts were performed between groups. Differentially expressed genes (DEGs) were identified using adjusted p-value < 0.05 (Benjamini–Hochberg FDR) and absolute log₂ fold change > 1.5.

Pathway and transcription factor analyses. Gene set enrichment analysis (GSEA) ^30^ was performed using clusterProfiler (v3.18.1) ^31^ with MSigDB Hallmark gene sets. Transcription factor analysis was performed using decoupleR (version 2.4.0)^32, 33^. Heatmaps were generated using pheatmap (v1.0.12) with z-score-transformed normalised counts, unsupervised hierarchical clustering (Euclidean distance, complete linkage), and group annotations. Volcano plots were generated using ggplot2 (v3.3.6) with gene labelling via ggrepel (v0.9.1).

### Re-analysis of publicly available human single-cell RNA-sequencing datasets

Publicly available scRNA-seq datasets from two independent human type-2 skin inflammatory conditions were retrieved from NCBI GEO: atopic dermatitis (GSE204762) and (GSE213849). CD4 T-cell clusters were identified based on positive expression of CD4 and CD3E together with absence of CD8A expression. For the atopic dermatitis dataset GSE204762, cells from lesional samples (excluding sample MGH106) and healthy controls were included in the analysis. To avoid patient overrepresentation, one sample from each patient and one sample from each healthy donor were selected for downstream analysis. For the GSE213849 dataset, all atopic dermatitis samples (n = 5) and all healthy control samples (n = 5) were included in the analysis. Raw count matrices were downloaded from the Gene Expression Omnibus (GEO) and analysed using the Seurat package (v4) in R. Raw 10X Genomics matrices were imported using the Read10X function and converted into Seurat objects using CreateSeuratObject with filtering thresholds of a minimum of 3 cells per gene and 200 detected features per cell. Metadata, including disease status and sex, were manually annotated for each sample prior to integration. Individual datasets were normalised using the NormalizeData function, and the top 2,000 highly variable genes were identified using FindVariableFeatures with the “vst” selection method. Integration anchors were identified using FindIntegrationAnchors, followed by dataset integration using IntegrateData to reduce batch effects across samples. The integrated assay was scaled using ScaleData, and dimensionality reduction was performed by principal component analysis (PCA) using the first 50 principal components. Uniform Manifold Approximation and Projection (UMAP) was subsequently performed using the first 30 principal components (n.neighbors = 15, min.dist = 0.1) for two-dimensional visualisation of cellular heterogeneity. Cell clustering was performed using the shared nearest neighbour (SNN) graph-based clustering approach implemented through the FindNeighbors and FindClusters functions in Seurat, with a clustering resolution of 0.1. Cluster-specific marker genes were identified using the FindAllMarkers function with Wilcoxon rank-sum testing, retaining positively enriched genes with a minimum expression in 25% of cells (min.pct = 0.25) and a log fold-change threshold of 0.25. Cell populations were annotated based on canonical lineage markers, and CD4 T-cell clusters were identified based on positive expression of CD4 and CD3E together with absence of CD8A expression. Differentially expressed genes (DEGs) between female and male CD4 T cells were identified using the FindMarkers function with Wilcoxon rank-sum testing (test.use = "wilcox"), using a minimum expression threshold of 10% of cells (min.pct = 0.1) and a log fold-change cutoff of 0.25. Additional DEG analyses were performed in GATA3-high CD4 T-cell subsets and individual CD4 T-cell clusters to investigate sex-specific transcriptional differences within distinct T-helper cell populations. Gene expression visualisation was performed using the FeaturePlot and DimPlot functions in Seurat.

#### Identification and purification of the Th2 cell population

To identify the Th2-equivalent cell population in each dataset, clusters were annotated using a combination of the Cluster Identity Predictor (CIPR) programme and marker gene expression from PangloDB. Th2 cells were defined by co-expression of CD4, CD3E, and GATA3, and clusters satisfying these criteria were selected for downstream analysis. Expression of IL13 was examined within the selected cluster(s) using FeaturePlot to confirm Th2 identity. To ensure transcriptional homogeneity and exclude any contaminating cytotoxic cells, cells expressing CD8A within the selected Th2 cluster(s) were identified and removed prior to differential expression analysis. Both male and female donors were included in the UMAP embeddings and visualised together.

#### Sex-stratified differential expression analysis

Within the purified CD4^+^CD3e^+^GATA3^+^ Th2 population, differentially expressed genes between female and male donors were identified using a pseudobulk approach. Donor identity was used as the replicate unit to generate aggregated count matrices per sex per donor. Differential expression analysis was performed using DESeq2 (v1.34.0) in R (v4.1.1), in which a log₂ fold change was computed for each gene and statistical significance was assessed using the Wald test. Genes with an adjusted p-value < 0.05 (Benjamini– Hochberg FDR) and an absolute log₂ fold change ≥ 1.0 were considered differentially expressed. Upregulated and downregulated gene sets were defined separately for each dataset.

#### Cross-species integration and Venn diagram analysis

Differentially expressed genes identified in the human datasets (female-upregulated and female-downregulated GATA3^+^ Th2 cells) were intersected with the corresponding upregulated and downregulated DEG sets from female murine Cyp11a1-knockout versus control Th2 cells generated in this study. Four-way Venn diagram analysis was performed independently for each human dataset (GSE213849 and GSE204762), incorporating the four gene sets: Cyp11a1-KO upregulated (murine), Cyp11a1-KO downregulated (murine), human female-upregulated, and human female-downregulated. Venn diagrams were constructed using InteractiVenn. The final consensus gene module was defined as the intersection of the three upregulated gene sets across all conditions: murine Cyp11a1-KO female Th2 cells, GSE213849 female GATA3+ T helper cells, and GSE204762 female GATA3+ T helper cells. Gene expression levels were visualised using heatmaps, violin plots, and bar plots generated with the pheatmap package (v1.0.12) and the dittoSeq R package (v1.2.4).

### Statistical analysis

For RNA-seq data, p-values were derived using the default methodologies in DESeq2 with Benjamini–Hochberg correction for multiple testing. For flow cytometry experiments, statistical significance was assessed using unpaired two-tailed Student’s t-test in GraphPad Prism 9 (GraphPad Software). A p-value < 0.05 was considered statistically significant. Statistical comparisons in figures are denoted as follows: *p < 0.05, **p < 0.01, ***p < 0.001, ****p < 0.0001; ns, not significant.

## Acknowledgements

We would like to thank Joana Cerveira, Cytometry facility, Dept. of Pathology and UBS animal facility, Gurdon Institute, for their technical help and animal husbandry. We used BioRender.com to generate the graphical illustrations presented in this manuscript.

## Author contribution

Conceptualisation: JP and BM. Methodology: JP, QZ, SC and CX. Investigation: JP, QZ and CX. Data analysis: JP, QZ, SC and CX. Data visualisation: JP, SC and QZ. Bioinformatics analyses: QZ, SC and CX. Writing the manuscript (original draft): JP and QZ. Editing the manuscript: SC, QZ, JP and BM. Supervision: BM. Project Administration: BM. Funding acquisition: BM. All authors read and approved the final manuscript.

## Data and materials availability

All newly generated RNA sequencing data used in this study are available at GEO with the accession number GSE330480 (password: wxgjcuichpibhan).

---

**Supplementary Figure S1. Comparing baseline proliferation and cytokine expression capacity of male and female Th2 cells.**

**(A)** Division index of control female (Ctrl_F) and control male (Ctrl_M) Th2 cells, taken from the experiments presented in Figure 4B and 4C. Male control Th2 cells divided slightly more than female control Th2 cells (*p < 0.05).

(B) Percentage of IL-4⁺ cells in Ctrl_F and Ctrl_M Th2 cells, from the experiments presented in Figure 4F (ns, p > 0.05).

(C) Percentage of IL-13⁺ cells in Ctrl_F and Ctrl_M Th2 cells, from the experiments presented in Figure 4E (ns, p > 0.05). All error bars indicate mean ± SEM. P values were calculated using two-tailed unpaired Student’s t test. Each dot represents one independent biological sample (n=6-11).

## REFERENCES

1 Walker JA, McKenzie ANJ. T(H)2 cell development and function. Nat Rev Immunol 2018; 18:121–133.

2 Paul WE, Zhu J. How are T(H)2-type immune responses initiated and amplified? Nat Rev Immunol 2010; 10:225–235.

3 Licona-Limon P, Kim LK, Palm NW, Flavell RA. TH2, allergy and group 2 innate lymphoid cells. Nat Immunol 2013; 14:536–542.

4 Akdis CA, Arkwright PD, Bruggen MC et al. Type 2 immunity in the skin and lungs. Allergy 2020; 75:1582–1605.

5 Shah R, Newcomb DC. Sex Bias in Asthma Prevalence and Pathogenesis. Front Immunol 2018; 9:2997.

6 Zein JG, Erzurum SC. Asthma is Different in Women. Curr Allergy Asthma Rep 2015; 15:28.

7 Zhang GQ, Ozuygur Ermis SS, Radinger M, Bossios A, Kankaanranta H, Nwaru B. Sex Disparities in Asthma Development and Clinical Outcomes: Implications for Treatment Strategies. J Asthma Allergy 2022; 15:231–247.

8 Hoffmann JP, Liu JA, Seddu K, Klein SL. Sex hormone signaling and regulation of immune function. Immunity 2023; 56:2472–2491.

9 Mahata B, Zhang X, Kolodziejczyk AA et al. Single-cell RNA sequencing reveals T helper cells synthesizing steroids de novo to contribute to immune homeostasis. Cell Rep 2014; 7:1130–1142.

10 Mahata B, Pramanik J, van der Weyden L et al. Tumors induce de novo steroid biosynthesis in T cells to evade immunity. Nat Commun 2020; 11:3588.

11 Wang M, Ramirez J, Han J et al. The steroidogenic enzyme Cyp11a1 is essential for development of peanut-induced intestinal anaphylaxis. J Allergy Clin Immunol 2013; 132:1174–1183 e1178.

12 Pramanik J, Zhao Q, Yamashita-Kanemaru Y, et al. Primitive steroidogenesis in mast cells: A novel regulatory mechanism for mast cell function. bioRxiv 2025:2025.2002.2005.636621.

13 Zhao Q, Pramanik J, Lu Y et al. Perturbing local steroidogenesis to improve breast cancer immunity. Nat Commun 2025; 16:3945.

14 Wells AD, Morawski PA. New roles for cyclin-dependent kinases in T cell biology: linking cell division and differentiation. Nat Rev Immunol 2014; 14:261–270.

15 Bird JJ, Brown DR, Mullen AC et al. Helper T cell differentiation is controlled by the cell cycle. Immunity 1998; 9:229–237.

16 Zhu J, Yamane H, Paul WE. Differentiation of effector CD4 T cell populations (*). Annu Rev Immunol 2010; 28:445–489.

17 Fiskin E, Eraslan G, Alora-Palli MB et al. Multi-modal skin atlas identifies a multicellular immune-stromal community associated with disrupted cornification and specific T cell expansion in atopic dermatitis. Nat Commun 2026; 17.

18 Calugareanu A, Specque F, Demouche S et al. Transcriptomic Landscape of Prurigo Nodularis Lesional Skin CD3+ T Cells Using Single-Cell RNA Sequencing. J Invest Dermatol 2023; 143:2525–2529 e2525.

19 Dou DR, Zhao Y, Belk JA et al. Xist ribonucleoproteins promote female sex-biased autoimmunity. Cell 2024; 187:733–749 e716.

20 Chang HY, Chung L, Davis MM et al. RNA origin of sex-biased immunity. Mol Ther Nucleic Acids 2026; 37:102853.

21 Welte T, Goulois A, Stadler MB et al. Convergence of multiple RNA-silencing pathways on GW182/TNRC6. Mol Cell 2023; 83:2478–2492 e2478.

22 Pramanik J, Shaji SK, Zaman M et al. Drug repurposing reveals posaconazole as a CYP11A1 inhibitor enhancing anti-tumor immunity. iScience 2025; 28:112488.

23 Acharya N, Madi A, Zhang H et al. Endogenous Glucocorticoid Signaling Regulates CD8(+) T Cell Differentiation and Development of Dysfunction in the Tumor Microenvironment. Immunity 2020; 53:658–671 e656.

24 Yang D, Huang L, Cheng H et al. A cell-intrinsic glucocorticoid biosynthesis and sensing circuit maintains a homeostatic Th17 cell state. Immunity 2026; 59:2232–2249 e2210.

25 Klein SL, Flanagan KL. Sex differences in immune responses. Nat Rev Immunol 2016; 16:626–638.

26 Laffont S, Blanquart E, Savignac M et al. Androgen signaling negatively controls group 2 innate lymphoid cells. J Exp Med 2017; 214:1581–1592.

27 Panten J, Del Prete S, Cleland JP et al. Four Core Genotypes mice harbour a 3.2MB X-Y translocation that perturbs Tlr7 dosage. Nat Commun 2024; 15:8814.

28 Souyris M, Cenac C, Azar P et al. TLR7 escapes X chromosome inactivation in immune cells. Sci Immunol 2018; 3.

29 Love MI, Huber W, Anders S. Moderated estimation of fold change and dispersion for RNA-seq data with DESeq2. Genome biology 2014; 15:1–21.

30 Subramanian A, Tamayo P, Mootha VK et al. Gene set enrichment analysis: a knowledge-based approach for interpreting genome-wide expression profiles. Proceedings of the National Academy of Sciences 2005; 102:15545–15550.

31 Yu G, Wang L-G, Han Y, He Q-Y. clusterProfiler: an R package for comparing biological themes among gene clusters. Omics: a journal of integrative biology 2012; 16:284–287.

32 Badia IMP, Velez Santiago J, Braunger J, et al. decoupleR: ensemble of computational methods to infer biological activities from omics data. Bioinform Adv 2022; 2:vbac016.

33 Garcia-Alonso L, Holland CH, Ibrahim MM, Turei D, Saez-Rodriguez J. Benchmark and integration of resources for the estimation of human transcription factor activities. Genome Res 2019; 29:1363–1375.

