## Supplementary Figure S1 for "A cell-intrinsic steroidogenic checkpoint constrains female Th2 effector output independently of gonadal hormones"

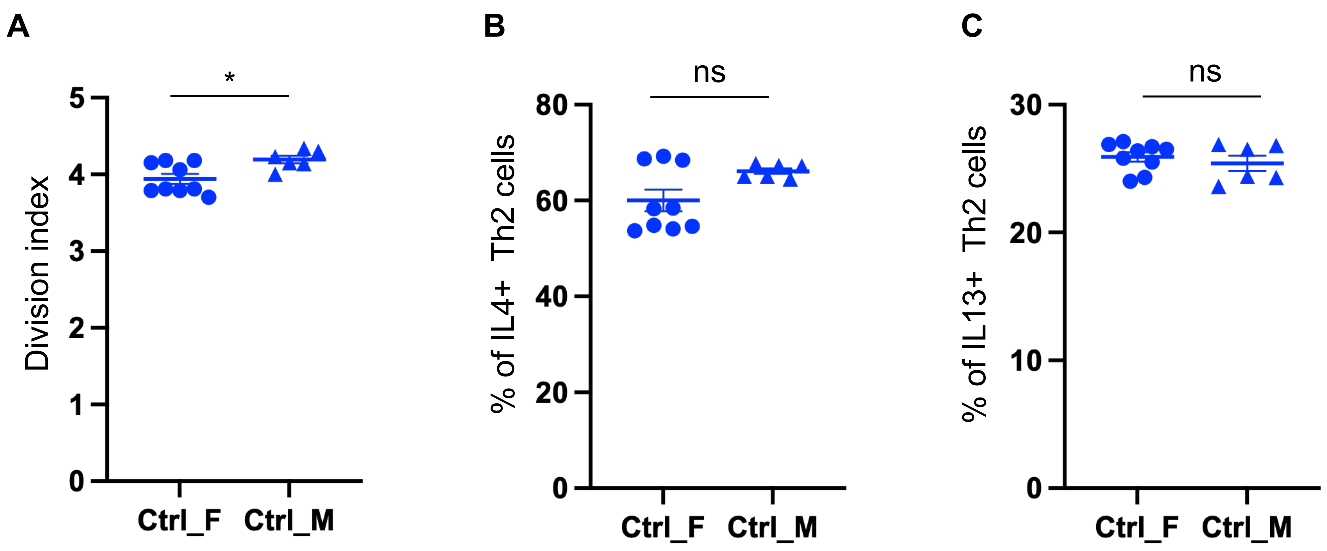


**Supplementary Figure S1. Comparing baseline proliferation and cytokine expression capacity of male and female Th2 cells.**
**(A)** Division index of control female (Ctrl_F) and control male (Ctrl_M) Th2 cells, taken from the experiments presented in Figure 4B and 4C. Male control Th2 cells divided slightly more than female control Th2 cells (*$p < 0.05$).
(B) Percentage of IL-$4^+$ cells in Ctrl_F and Ctrl_M Th2 cells, from the experiments presented in Figure 4F (ns, $p > 0.05$).
(C) Percentage of IL-$13^+$ cells in Ctrl_F and Ctrl_M Th2 cells, from the experiments presented in Figure 4E (ns, $p > 0.05$). All error bars indicate mean ± SEM. P values were calculated using  two-tailed unpaired Student’s t test. Each dot represents one independent biological sample (n=6-11).
